# Relaxed selection during a recent human expansion

**DOI:** 10.1101/064691

**Authors:** S. Peischl, I. Dupanloup, A. Foucal, M. Jomphe, V. Bruat, J.-C. Grenier, A. Gouy, E. Gbeha, L. Bosshard, E. Hip-Ki, M. Agbessi, A. Hodgkinson, H. Vézina, P. Awadalla, L. Excoffier

## Abstract

Humans have colonized the planet through a series of range expansions, which deeply impacted genetic diversity in newly settled areas and potentially increased the frequency of deleterious mutations on expanding wave fronts. To test this prediction, we studied the genomic diversity of French Canadians who colonized Quebec in the 17^th^ century. We used historical information and records from ∼4000 ascending genealogies to select individuals whose ancestors lived mostly on the colonizing wave front and individuals whose ancestors remained in the core of the settlement. Comparison of exomic diversity reveals that i) both new and low frequency variants are significantly more deleterious in front than in core individuals, ii) equally deleterious mutations are at higher frequencies in front individuals, and iii) front individuals are two times more likely to be homozygous for rare very deleterious mutations present in Europeans. These differences have emerged in the past 6-9 generations and cannot be explained by differential inbreeding, but are consistent with relaxed selection on the wave front. Modeling the evolution of rare variants allowed us to estimate their associated selection coefficients as well as front and core effective sizes. Even though range expansions had a limited impact on the overall fitness of French Canadians, they could explain the higher prevalence of recessive genetic diseases in recently settled regions. Since we show that modern human populations are experiencing differential strength of purifying selection, similar processes might have happened throughout human history, contributing to a higher mutation load in populations that have undergone spatial expansions.

## Introduction

The impact of recent demographic changes or single bottlenecks on the overall fitness of populations is still highly debated (Lohmueller et al. 2008; Lohmueller 2014; Simons et al. 2014; Do et al. 2015; Gravel 2016), but simulation and theoretical approaches suggest that populations on expanding wave fronts accumulate deleterious mutations over time (Peischl et al. 2013; Peischl et al. 2015), and thus build-up an expansion load (Peischl et al. 2013). This accumulation is mainly driven by low population densities and strong genetic drift at the wave front promoting genetic surfing of neutral and selected variants (Peischl et al. 2013). This relatively inefficient selection on the wave front leads to the preservation of many new mutations, unless very deleterious (Peischl et al. 2013). After a range expansion, both a decrease of diversity and an increase in the recessive mutation load with distance from the source is expected (Kirkpatrick and Jarne 2000; Peischl and Excoffier 2015). This pattern has recently been shown to occur in non-African human populations, where a gradient of recessive load has been observed between North Africa and the Americas (Henn et al. 2015b). Whereas the bottleneck out of Africa that started about 50 Kya (e.g. Gravel et al. 2011) must have created a mutation load, the exact dynamics of this load increase due to the expansion process is still unknown. It is also unclear if a much more recent expansion could have had a significant impact on the genetic load of populations.

The settlement of Quebec can be considered as a series of demographic and spatial expansions following initial bottlenecks. Indeed, the majority of the 6.5 million French Canadians living in Quebec are the descendants of about 8,500 founder immigrants of mostly French origin (Charbonneau et al. 2000; Laberge et al. 2005). This French immigration started with the founding of a few settlements along the Saint-Lawrence river at the beginning of the 17^th^ century (Charbonneau et al. 2000). Most new settlements were restricted to the Saint-Lawrence valley until the 19^th^ century, after which new remote territories began to be colonized. Bottlenecks and serial founder effects occurring during range expansions are thought to have profoundly affected patterns of genetic diversity, leading to large frequency differences when compared to the French source population (Laberge et al. 2005). Even though the French Canadian population has expanded 700 fold in about 300 years, its genetic diversity is actually not what is expected in a single panmictic, exponentially growing, population, as allele frequencies have drifted much more than expected in a fast growing population (Heyer 1995; Heyer 1999). Indeed, it has been shown that genetic surfing (Klopfstein et al. 2006; Peischl et al. 2016) has occurred during the recent colonization of Saguenay-Lac St-Jean area (Moreau et al. 2011), and that the fertility of women on the wave front was 25% higher than those living in the core of the settlement, giving them more opportunity to transmit their genes to later generations. In addition, female fertility was found to be heritable on the front but not on the core (Moreau et al. 2011), a property that further contributes to lower the effective size of the population (Austerlitz and Heyer 1998; Sibert et al. 2002) and to enhance drift on the wave front. Social transmission of fertility (Austerlitz and Heyer 1998) and genetic surfing during range expansions or a combination of both (Moreau et al. 2011) have been proposed to explain a rapid increase of some low frequency variants. It thus seems that differences in allele frequencies between French Canadians and continental Europe are due to a mixture of the random sampling of initial immigrants (founder effect) and of strong genetic drift having occurred in Quebec after the initial settlement, resulting in a genetically and geographically structured population of French Canadians (Bherer et al. 2011).

The demographic history of Quebec has not only affected patterns of neutral diversity, but also the prevalence of some genetic diseases independently from inbreeding (De Braekeleer 1991; Heyer 1995; Laberge et al. 2005; Yotova et al. 2005), as well as the average selective effect of segregating variants (Casals et al. 2013). Even though French Canadians have fewer mutations segregating in the population than the French, these mutations are found at loci which are, on average, much more conserved, and thus are potentially more deleterious than those segregating in the French population (Casals et al. 2013). Recurrent founder effects, low densities and intergenerational correlation in reproductive success could all contribute to increase drift and reduce the efficacy of selection on expanding wave fronts, and thus lead to the development of a stronger mutation load (Gravel 2016). It is therefore likely that the excess of low frequency deleterious variants observed in French Canadian individuals (e.g. Casals et al. 2013), could be at least partly due to the expansion process rather than to the sole initial bottleneck.

To better understand and quantify the effect of a recent expansion process on the amount and pattern of mutation load, we screened the ascending genealogies of 3916 individuals from the CARTaGENE cohort (Awadalla et al. 2013) that were linked to the BALSAC genealogical database (http://balsac.uqac.ca/). Using stringent criteria on the quality of genealogical information (see Methods), we selected 51 (front) individuals whose ancestors were as close as possible to the front of the colonization of Quebec, and 51 (core) individuals whose ancestors were as far as possible from the front (see Methods, **Fig. 1**, and **Supporting Animation S1 and S2**). We then sequenced these 102 individuals at very high coverage (mean 89.5X, range 67X-128X) for ∼106.5 Mb of exomic and UTR regions and contrasted their genomic diversity to detect if sites with various degrees of conservation and deleteriousness had been differentially impacted by selection.

**Figure 1:**
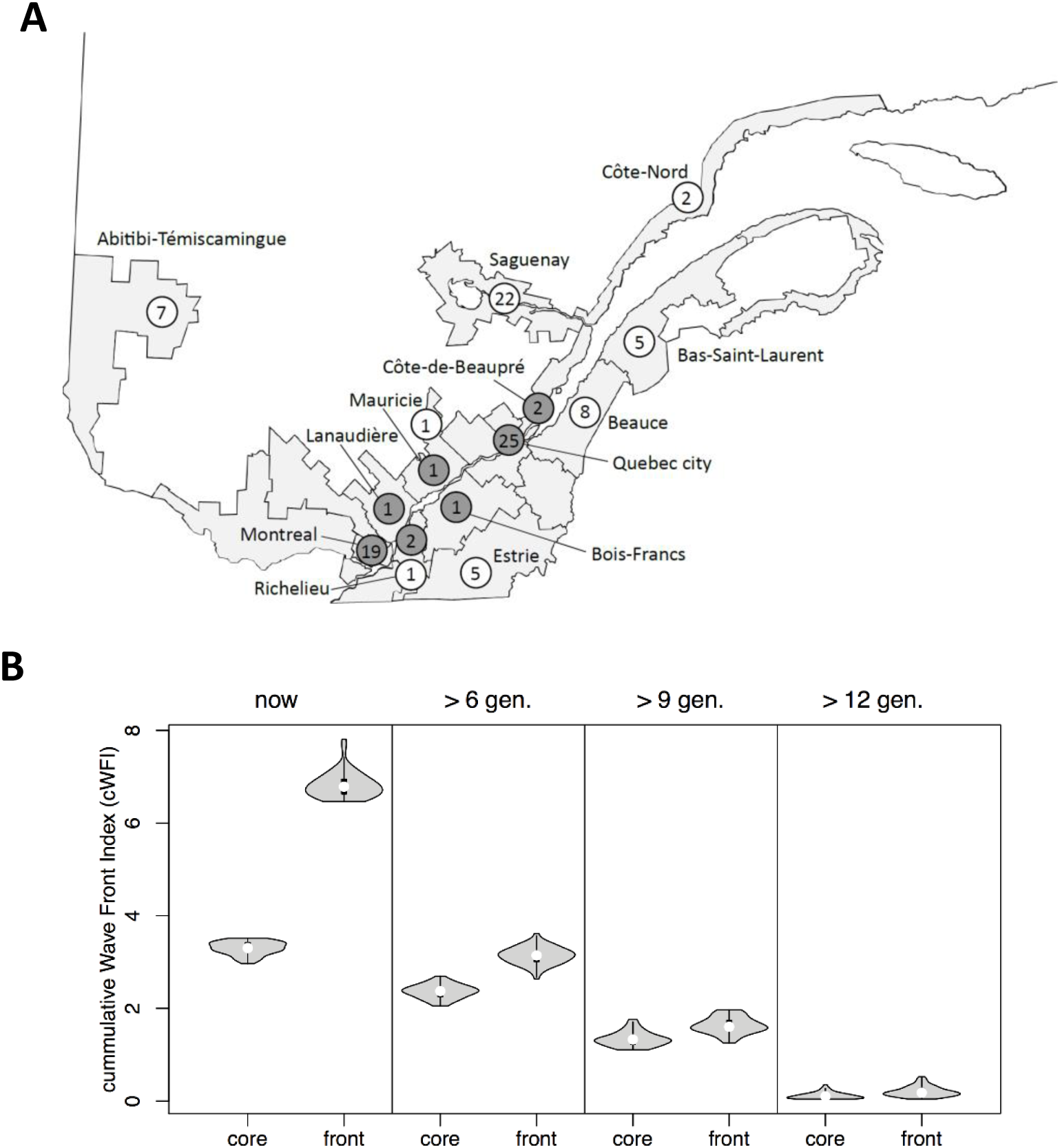
Location and number of sampled individuals and distribution of the cumulative Wave front Indices (cWFI). **A**: Front and core sampled individuals are shown in white and gray, respectively. The numbers inside circles indicate the sample size for each location. **B**: The leftmost panel shows the distribution of cWFI among sampled individuals. The other three panels display the cWFI of the ancestors of the sampled individuals that lived 6, 9 or 12 generations ago, which shows that observed differences in cWFI between current samples have mostly emerged in the 6 most recent generations.

## Results

### French Canadians vs. Europeans

French Canadians are genetically very divergent from three European populations of the 1000 Genome phase 3 panel (The Genomes Project 2015) (Great Britain, Spain, and Italy, **Supplementary Fig. S1**), as expected after a strong bottleneck. When focusing on SNPs shared between French Canadians and Europeans and thus on relatively high frequency variants, core individuals are found genetically closer to European samples than front individuals (**Supplementary Fig. S1B**), in keeping with stronger drift having occurred on the wave front. If we assess the functional impact of point mutations with GERP Rejected Substitution (GERP-RS) scores (Davydov et al. 2010), sites polymorphic in French Canadians are on average more conserved than sites polymorphic in European. Thus even though French Canadians have fewer polymorphic sites than 1000G populations from Europe, their variants are on average potentially more deleterious than those found in European samples (**Fig. 2A**), in line with previous results (Casals et al. 2013). Note that this results still holds if we focus only on SNPs that are shared between 1000G and Quebec samples, even though the distributions are slightly more overlapping (**Fig 2A**).

**Figure 2:**
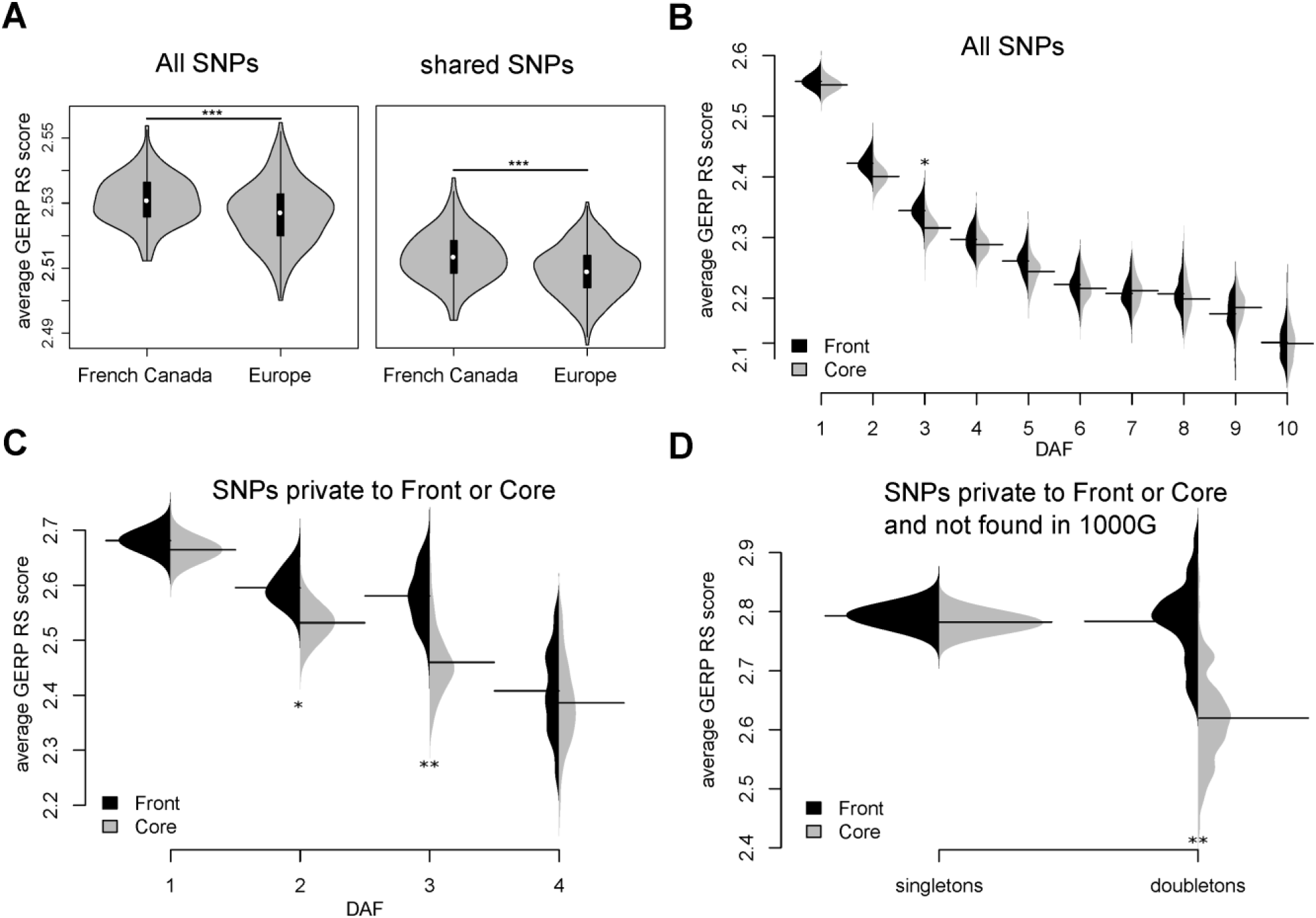
**A**: Distributions of average GERP-RS scores per site per individual in three European 1000G populations, as well as in core and front individuals. Left: All sites. Right: Sites shared between 1000G samples and Quebec (t-test p-values = 10^−7^ and 10^−5^, respectively). **B**: Average GERP score per site having different Derived Allele Frequencies (DAF). The solid horizontal lines show the average GERP RS score per site. The violinplots show the the average GERP score distribution obtained by bootstrap (1000 replicates). **C:** Like B, but for mutations private to the front or to the core. **D**: Like B but for singletons and doubletons that are private to front or core and not found in the 1000G phase 3 panel. For the sake of clarity, higher DAF classes are not shown in panels B- D. Only SNPs with GERP scores larger than 0 were used for the calculations of GERP scores in all panels. Asterisks indicate signifcance levels obtained by permutation tests: * *p* < 0.05, ** *p* < 0.01, *** *p* < 0.001.

### Genomic diversity of front and core individuals

In French Canadians, front individuals have a significantly smaller number of variants than core individuals (**Table 1**) consistent with higher rates of drift. The allele frequencies in front and core individuals are overall very similar (**Supplementary Fig. S3**), but there is a significant deficit of singletons on the front as compared to the core (p_perm_ < 0.001, **Supplementary Table S4, Supplementary Fig. S4**), which is balanced by an excess of doubletons on the front (p_perm_ < 0.001). Note that this pattern is consistently found for all GERP-RS score categories (**Supplementary Figs. S4**-**S7** and **S9**). We then looked whether genes containing SNPs with large frequency differences between front and core (i.e. those with FST p-value < 0.01) were overly represented in some gene ontology (GO) categories. The top 25 significantly enriched GO categories (**Supplementary Table S1**) are generally involved in very conserved processes like gene expression, development and cell growth (**Supplementary Fig. S15**), suggestive of a relaxation of selection rather than specific adaptations to the front environment.

**Table 1:** Summary of genetic diversity in front and core samples.

| Type and number of polymorphism | core<br>(n=51) |  | front<br>(n=51) | Total<br>(n=102) |
| --- | --- | --- | --- | --- |
| Total No. of SNPs |  |  |  | 426,301 |
| No. of SNPs with inferred ancestral/derived state | 314,483 | > | 308,396 | 396,424 |
| No. of SNPs without missing data | 266,547 | > | 261,355 | 328,372 |
| No. of exonic SNP | 83,653 | > | 81,763 | 107,525 |
| No. of non-synonymous SNP | 40,750 | > | 39,595 | 55,133 |
| No. of SNPs private to one of the two groups of individuals | 78,310 | > | 72,353 | 150,663 |
| No. of SNPs without missing data and not seen in 1000G phase 3 panel | 31,608 | > | 29,811 | 56,669 |
| No. of SNPs without missing data, private to one of the two groups, and not seen in 1000G phase 3 panel | 26,858 | > | 25,061 | 51,919 |
| No. of indels | 33,789 | > | 33,297 | 43,081 |
| Heterozygosity |  |  |  |  |
| All sites | 0.0588 | ≈ | 0.0586 |  |
| Exons | 0.0548 | ≈ | 0.0547 |  |
| Introns | 0.0632 | ≈ | 0.0630 |  |
| 5' UTR | 0.0489 | ≈ | 0.0487 |  |
| 3'UTR | 0.0623 | ≈ | 0.0623 |  |

Significant differences between front and core are indicated by “>” (permutation test, $p_{\text{perm}} < 0.001$ ), and non-significant differences are indicated by “≈” ( $p > 0.05$ ).

### Low frequency variants in front individuals are more conserved

The examination of low frequency variants that are enriched for deleterious mutations (Boyko et al. 2008; Nelson et al. 2012; Kiezun et al. 2013) should allow us to better evidence the presence of differential selection between front and core. We indeed find a negative relationship between the frequency of mutations and their average GERP-RS scores (**Fig. 2B**), and low frequency variants (DAF < 5%) have significantly larger GERP-RS scores, and are thus potentially more deleterious, on the front than in the core (p_perm_ = 0.038). Since new variants should also be enriched for deleterious mutations (Boyko et al. 2008; Keinan and Clark 2012), we then focused on mutations private to front or to core individuals. With this additional filtering, the differences in GERP-RS scores between front and core for low frequency mutations are much more pronounced (**Fig. 2C**), with significant differences for both doubletons and tripletons (p_perm_ = 0.03 and p_perm_ = 0.0025, respectively). We checked that these results were not due to our use of the GERP-RS scoring system by repeating analyses using CADD conservation scores (Kircher et al. 2014). We find overall very similar evidence of reduced selection in front populations (**Supplementary Figs. S8,** and **S10–S13**) for point mutations and for indels identified as under selection by CADD, suggesting that our results are robust to alternative deleteriousness scoring systems.

### New deleterious mutations have reached higher frequencies on the front

We further enriched our data for new mutations that occurred during the colonization of Quebec by focusing only on French Canadian mutations that are not observed in the entire 1000G phase 3 panel and are private either to the core or to the front samples. In this filtered data set, we find a significant excess of predicted deleterious (GERP-RS score > 2) singletons in the core (p_perm_ < 0.001), and an excess of doubletons in the front (p_perm_ < 0.001, **Supplementary Table S4**). Interestingly, the doubletons on the front are as conserved as singletons in both core and front samples, suggesting that doubletons on the front are variants that would be singletons in the core (**Fig. 2D**). To see if inbreeding could explain the observed excess of deleterious doubletons in the front, we compared samples from the region of Saguenay, where remote inbreeding is higher than in the rest of Quebec (**Supplementary Fig. S12**), with front samples coming from other regions of Quebec. We find that doubletons in less inbred non-Saguenay individuals are at loci that are on average more conserved than those of Saguenay individuals (**Supplementary Fig. S14**), showing that inbreeding cannot explain the increase in frequency of rare deleterious variants. Because it is difficult to estimate mutation load from sequence data (Lohmueller 2014), we then used the sum of GERP-RS scores of new or rare deleterious doubletons per individual across the four GERP-RS score categories as a proxy for mutation load. As shown in **Figure 3**, the cumulative GERP-RS scores are similar in front and core individuals for neutral sites (−2 < GERP-RS < 2), but significantly larger in front individuals for non-neutral GERP-RS score categories (GERP-RS ≥2), suggesting that differential selection has allowed mutations at more conserved sites to increase in frequency on the front.

**Figure 3:**
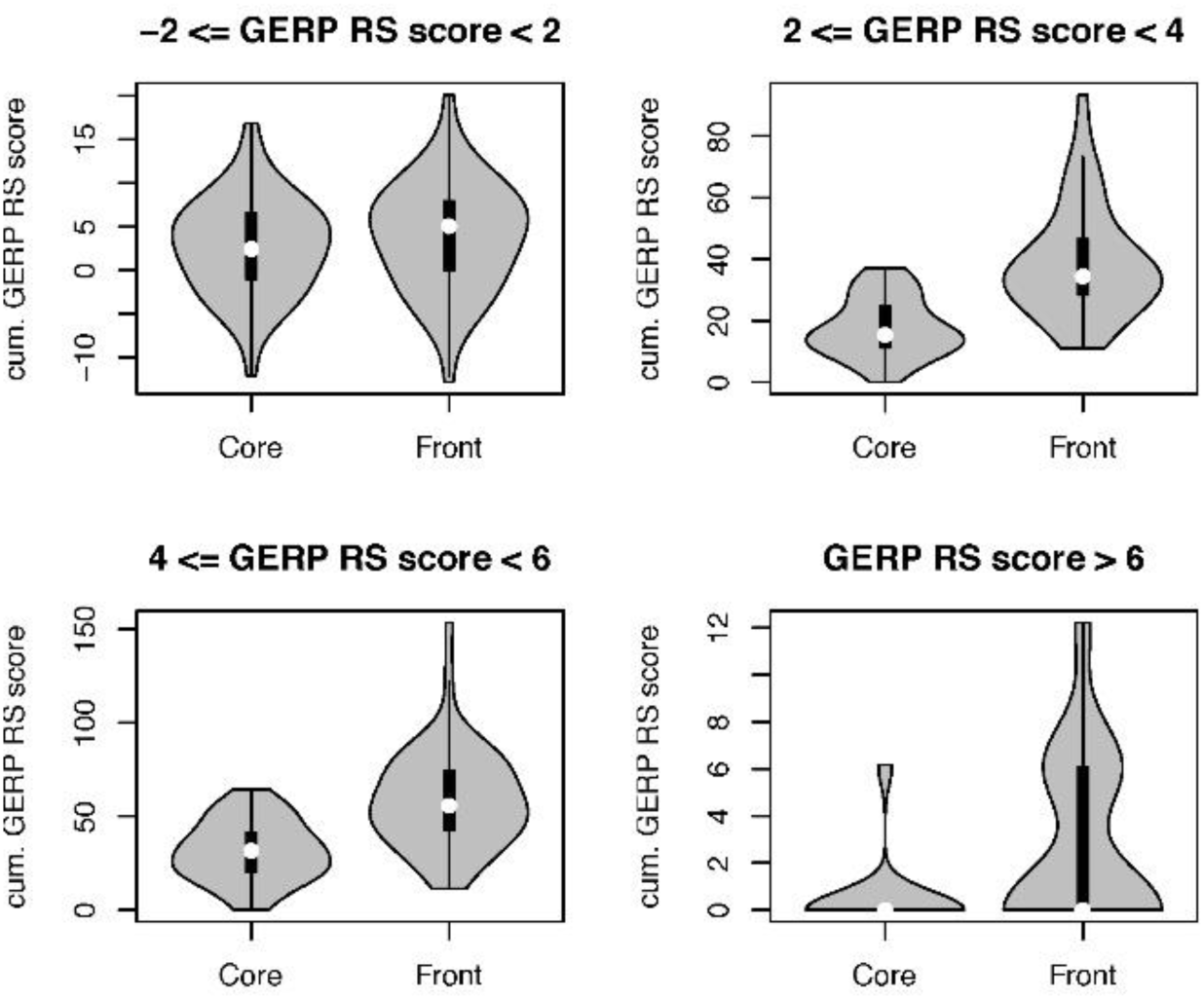
Distribution of the cumulative additive GERP-RS scores of doubletons in front and core individuals for different GERP-RS categories. Sites were considered if they were not seen in derived states in 1000G samples and if they were private to the core or to the front. Differences between front and core are significant for the three categories of sites potentially under selection (p = 10^−11^, 10^−9^, 10^−4^ for mildly, strongly, and extremely deleterious sites, respectively), but not for the neutral sites (−2 < GERP-RS score < 2, p = 0.34).

### Variants with low frequency in Europe have been more impacted by selection in the core

Because neutral sites should only be affected by drift and not by selection, stronger drift at the front should increase the variance of neutral allele frequencies (Gravel 2016), but should not affect their average frequency. In contrast, the frequency of deleterious variants should be smaller in the core if the purging of deleterious variants was more efficient. To test these predictions, we followed mutations that are singletons in European 1000G populations and that are still seen in Quebec. In agreement with theory, we find no significant difference in the average derived allele frequencies (*x_d_*) of European singletons predicted to be neutral (GERP-RS score between −2 and 2) (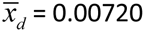 vs 0.00717 in front and core, respectively, p_perm_ = 0.34), and a slightly larger variance of derived allele frequencies on the front (s. d. (*x_d_*): 0.0163 vs 0.0159, p_perm_ = 0.072). Contrastingly, predicted deleterious sites have significant higher derived allele frequencies on the front than in the core (p_perm_ = 0.0146 for sites with GERP-RS score > 4), in keeping with higher selective pressures in the ancestry of core individuals.

Since differences between core and front individuals are strongest for rare alleles, these differences may have an impact on the homozygosity of recessive deleterious alleles and thus influence disease incidence. We used the ClinVar database (Landrum et al. 2014) to identify pathogenic variants (causing Mendelian disorders, Richards et al. 2015) in the set of SNPs segregating in French Canadians. The distribution of GERP RS scores for pathogenic variants is clearly shifted towards higher GERP RS scores as compared to the distribution for all SNPs loci (**Supplementary Figs. S23** and **S24**), confirming that GERP RS is a valid deleteriousness scoring system. We find that front individuals have a 11.8% higher probability to be homozygotes for these pathogenic variants than core individuals, suggesting that the expansion process has also affected disease causing mutations. For rare deleterious variants (i.e., derived singletons in Europe with GERP-RS score >2), this excess in homozygosity is 9.5 %. Of importance, this excess increases with GERP-RS scores and reaches approximately 90% (p_perm_ = 0.021) for sites with a GERP-RS score larger than 6 (**Fig. 4**). Note that this increase cannot be explained by the higher inbreeding level prevailing on the front, and that the differences in homozygosities between front and core become even more pronounced if one removes more inbred Saguenay individuals (p_perm_ = 0.008, **Fig. 4**). This last result shows that stronger purifying selection in the core rather than higher inbreeding on the front is directly responsible for the lower frequencies of deleterious mutations in the core.

**Figure 4:**
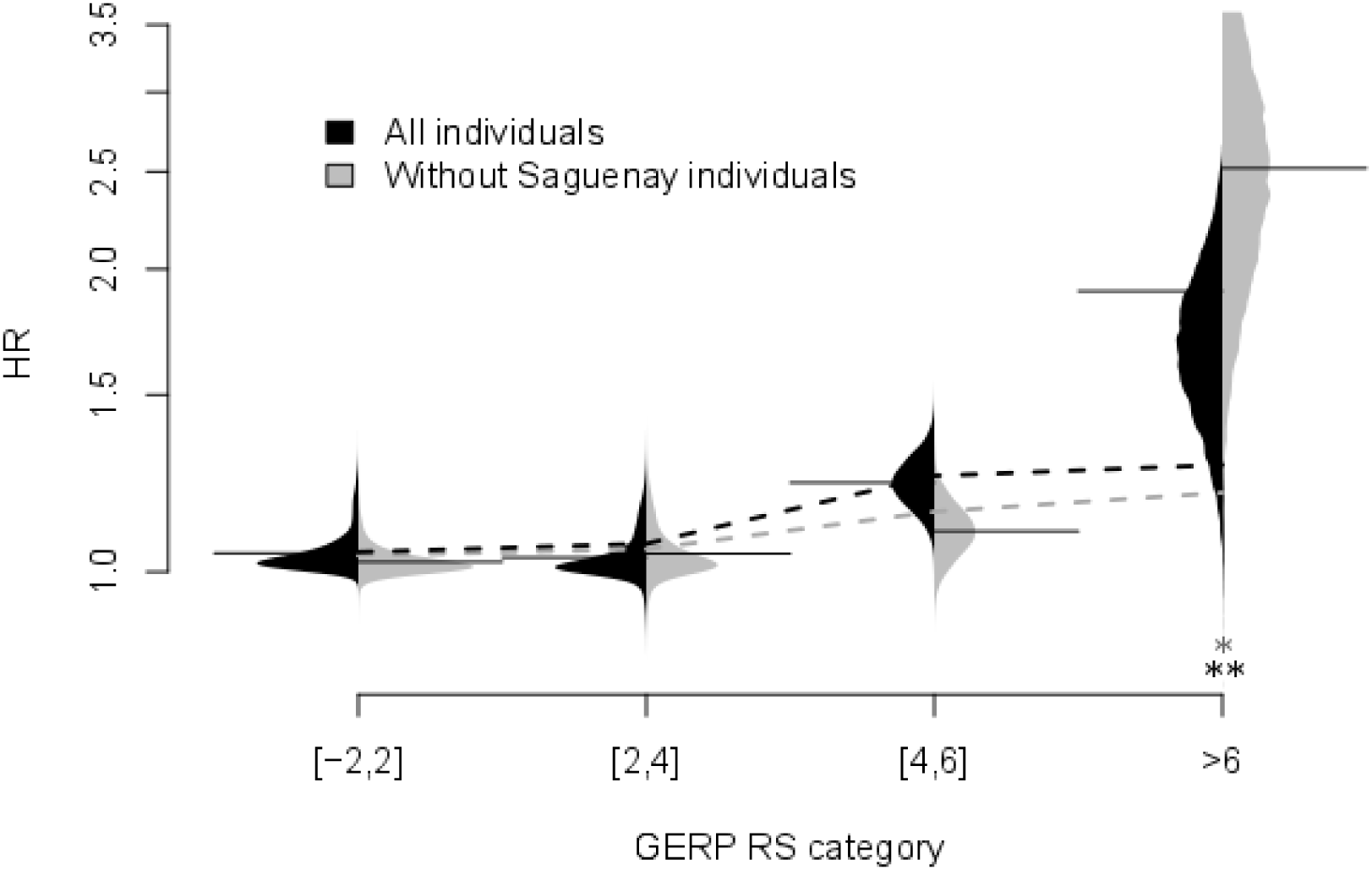
Ratio of expected homozygosity for variants that are singletons in European 1000G populations. 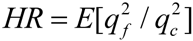 where *q_f_* and *q_c_* are derived allele frequencies in front and core individuals, respectively. The horizontal solid lines indicate HR for different GERP RS score categories. The dashed lines indicates the expected HR values that would be due to differences in estimated inbreeding levels between front and core, calculated as 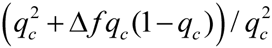, where Δ*f* = *f_front_* − *f_core_*. Violin plots show the distribution of 5000 bootstrap replicates. We find signifcant differences between the expected values for GERP RS scores > 6 (all individuals: *p* = 0.021, without Sageunay individuals: *p*= 0.008, obtained by bootstrap).

### Likelihood-based demographic and selection coefficient inferences

We used the allele frequency distributions of mutations that are singletons in European 1000G populations and that are still seen in Quebec to estimate the parameters of a simple demographic model for the settlement of French Canada. In this model, a small founding population splits off from the ancestral population, and then further splits into two subpopulations; the front and the core (**Fig. 5A**). We estimate the effective population size of the founding population (*N_BN_*), the front (*N_F_*), and the core (*N_C_*) under a maximum-likelihood framework based on inter-generational allele frequency transition matrices (see Methods for details). We report here results for a model in which we fix the duration of initial bottleneck to one generation, but the analysis of a model with a 7 generation bottleneck yields qualitatively similar results, which can be found in the Supporting Information (**Supplementary Fig. S25**). We infer that French Canadians passed through a bottleneck equivalent to 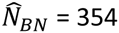 effective diploid individuals, and that the front population was about 2.5 smaller 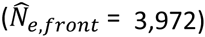 than the core population 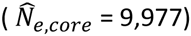 (**Fig. 5B**). We then used these maximum likelihood estimates (MLE) to estimate the contribution of the range expansion to the total variance in allele frequencies on the front as V_F_ = V_*BN*_ + V_*EXP*_, where V_BN_ is the variance in allele frequencies after the bottleneck, and V_EXP_ is the remaining variance due to the expansion process. We find that V_*EXP*_ explains about 20% of the total variance in allele frequencies that occurred since the initial settlement at the expansion front. Therefore, we estimate that under our simple model, 20% of the genetic divergence between Europe and the front has been generated by the expansion process, whereas the remaining 80% is due to the initial bottleneck shared by the core. We also estimated the strength of selection associated to rare variants under our estimated demographic model. In agreement with predictions, the MLE for the selection coefficient associated to predicted neutral variants is centered around zero, whereas the selection coefficients associated to predicted deleterious sites are clearly negative and decrease with increasing GERP RS score (**Fig. 6B**, maximum likelihood estimates and 95% confidence intervals: −0.006 < *ŝ*_GERP_[−2,2] = 0 < 0.006, −0.034 < *ŝ*_GERP_[2,4] = −0.024 < −0.013, −0.042 < *ŝ*_GERP_[4,6] = −0.032 < −0.022, −0.145 < *ŝ*_GERP_>6 = −0.072 < 0.001). Note that the most negative selection coefficient for GERP-RS > 6 is not significantly different from zero due to the small number of sites belonging to this category.

**Figure 5.**
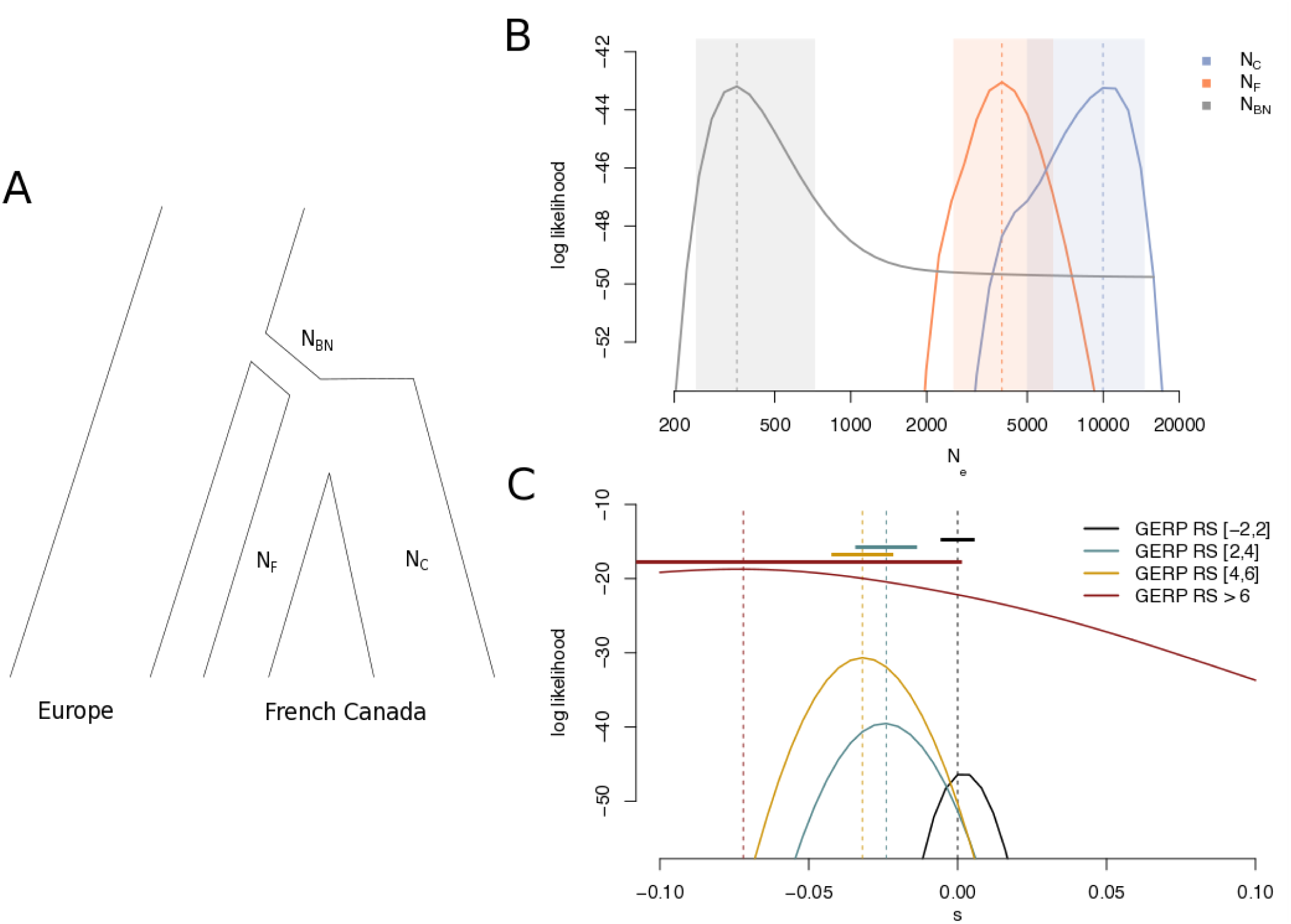
**A:** Sketch of the model used for maximum likelihood estimation. Likelihoods were calculated based on the expectation of the change in allele frequency distribution of rare variants (that is, singletons in the European sample). Marginal likelihoods and MLE for effective population sizes of bottleneck, and in front and core (**B**), and selection coeffcients for different GERP-RS categories (**C**). Shaded areas indicate 95% confidence intervals in (**B**), and horizonal bars indicate 95% confidence intervals in (**C**).

### Simulations can reproduce observed differences between front and core

Whereas it seems difficult to perform demographic inferences under a complex spatially explicit model, we can use forward simulations to see how well a model of range expansion can explain our observations (see Methods for details on the simulations). Our simulations reveal that the observed excess of singletons in core populations as well as the excess of doubletons in front populations are consistent with a model of range expansion (**Supplementary Fig. S19**), in keeping with previous results showing that range expansions leads to a flattening of the SFS (Sousa et al. 2014). Importantly, simulations also confirm these features of the SFS for negatively selected mutations (**Supplementary Fig. S19**). Our simulations also confirm that an excess of homozygosity should develop on the front and that it should increase with the deleteriousness of mutations (**Supplementary Fig. S20**), in keeping with the observed patterns in Quebec (**Fig. 4**). Together, these results show that a model of range expansion can well explain most of the observed differences between front and core populations in Quebec.

## Discusssion

The interaction between demography and selection has been a central theme in population genetics. A particularly hotly debated topic is whether and to what extent recent demography has affected the efficacy of selection in modern humans (Lohmueller et al. 2008; Lohmueller 2014; Simons et al. 2014; Do et al. 2015; Gravel 2016). The original conclusion that European population show a larger proportion of predicted deleterious variants when compared to African populations (Lohmueller et al. 2008) has been recently revisited in a series of studies that reached different and apparently opposite conclusions (reviewed in Lohmueller 2014). However, this controversy might have arisen because different studies focused on different patterns or processes. First, people focused either on measures of the efficacy of selection (the amount of change in load per generation) or on measures of the mutation load (see e.g. Gravel 2016, for a detailed study of this distinction). Second, people either measured the load as being due to co-dominant (Simons et al. 2014; Do et al. 2015) or partially recessive (Henn et al. 2015b) mutations, which can lead to drastically different conclusions about the consequences of demographic change on mutation load (Henn et al. 2015a; Henn et al. 2015b). Finally, most theoretical and empirical work has focused on the effects of bottlenecks and recent population growth, but ignored the out of Africa expansion process and the spatial structure of human populations (Sousa et al. 2014). While it has now been shown that the out of Africa expansions that started more than 50 kya have led to the buildup of a mutation load in non-Africans that is proportional to their distance from Africa (Henn et al. 2015b), it was unclear whether an expansion load could develop during much shorter expansions, if it could be evidenced in very recent or ongoing expansions, and what are the exact genomic signatures of this expansion load.

We have used here a unique combination of historical records, detailed genealogical information, and genomic data to study the impact of such a recent range expansion on functional genetic diversity, and to disentangle the effects of genetic drift, purifying selection, and inbreeding during an expansion. The significant differences we have detected between front and core individuals all suggest that relaxed purifying selection on the front slightly but rapidly increases the frequency of deleterious mutations. The fact that front and core individuals mainly diverged six generations ago with respect to the position of their ancestors to the colonization front (**Fig. 1B**) suggests that the relaxation of natural selection can affect remarkably quickly modern populations. The recent divergence between front and core populations (around 1780, **Supplementary Fig. S21**) has left traces in the genomic diversity of French Canadians that are of two kinds. First, front individuals show increased genetic drift relative to core individuals, as attested by their overall lower levels of diversity (**Table 1**), their larger genetic divergence from Europeans (**Supplementary Fig. S1**), and their lower estimated effective size (**Fig. 5B**). This result confirms the genetic surfing effect previously identified in the Saguenay Lac St-Jean region (Moreau et al. 2011), but it is not driven by samples from the Saguenay area (e.g. **Supplementary Fig. S14**). Rather, it is a property shared by all individuals with ancestors having lived on the front, and presently found in the most peripheral regions of Quebec (**Fig. 1**). Second, we find several lines of evidence showing relaxed selection in front individuals as compared to core individuals, which leads to the increase in frequency of rare and potentially deleterious variants. The evidence comes from the fact that sites targeted by mutations tend to be more conserved in front than in core individuals (**Fig. 2B-2D**), and that rare, putatively deleterious derived alleles, have a higher probability to be homozygous at the front (**Fig. 4**). Relaxed selection is especially obvious when one considers deleterious mutations that were at low frequencies (singletons) in Europe and that have been kept at lower frequencies in core than in front individuals, or mutations that are now at low frequencies in Quebec and that are occurring at more conserved sites (and thus potentially more deleterious) in front than in core individuals (e.g., private doubletons and tripletons in **Figs. 2C and 2D**).

At first sight, the increased frequency of rare and potentially deleterious alleles (i.e. doubletons) in front individuals could be attributed to their higher inbreeding levels. However, there are several lines of arguments against this interpretation. First, we note that there are about 5% more doubletons on the front than in the core (21,332 vs. 20,284, **Supplementary Fig. S4**), which cannot be explained by a difference in inbreeding level of only 0.3% (**Supplementary Fig. S2**). Instead, individual based simulations show that the excess of doubletons at the front is consistent with a model of range expansion (**Supplementary Figs. S4 and S19**). Second, the proportion of doubleton sites where both derived alleles are in the same individuals is smaller than expected (1/101=0.99%) in both front (0.651%) and core (0.646%) individuals, which is indicative of similar (p_perm_ = 0.898) levels of selection against derived homozygotes in both samples. Third, if higher inbreeding (and not relaxed selection) on the front had increased the frequency of all rare mutations irrespective of their deleterious effect, more deleterious mutations should have been better purged by selection than less deleterious mutations, and observed doubletons on the front should be on average less conserved. However, we find the opposite, with doubletons at the front being more conserved than in the core (**Fig. 2**), which means that the number of doubletons at highly conserved sites has increased proportionally more than at neutral sites. Fourth, we find that less inbred individuals from the front tend to have rare variants that are more deleterious than more inbred individuals from the Saguenay area (**Supplementary Figs. S2 and S14**). Finally, the difference in inbreeding level between front and core individuals cannot explain the 2-fold increased expected homozygosity for extremely deleterious variants on the front (**Fig. 4**), and removing Saguenay individuals from the analysis amplifies the excess of derived homozygotes on the front (**Fig. 4**). A model of range expansion can however explain the increase in derived homozygosity at the expansion front (**Supplementary Fig. S20**). Taken together, these results suggest that differences between front and core individuals are mainly driven by increased drift at the expansion front and more efficient selection against deleterious mutations in the core.

In line with previous results (Casals et al. 2013), we find that all French Canadians present a much larger mutation load than Europeans (**Fig. 2A**). Even though it has been proposed that this is the result of a mere founder effect (Casals et al. 2013), current French Canadians descend from ∼8500 French founders (Laberge et al. 2005), which implies a relatively mild founder effect that would take hundred to thousand generations to increase load to such an extent (Lohmueller et al. 2008; Peischl et al. 2013). More likely, this load could have been created during the initial settlement and range expansion that occurred in Quebec along the Saint-Lawrence valley. A major loss of diversity and an increase in the frequency of rare deleterious variants might indeed have occurred during the first 9 generations of the settlement of Quebec, until the middle of the 18^th^ century, before current front and core individuals actually diverged (**Fig. 1B**). The importance of these early generations is supported by genealogical analyses of the genetic contributions of the founders having lived at different periods. Early settlers have indeed contributed between 45% to 90% to the current French Canadian gene pool (Heyer 1995; Bherer et al. 2011), depending on the regions of Quebec, and early founders contributed proportionally more than later individuals to the current French Canadian gene pool (Heyer 1995; Bherer et al. 2011; Moreau et al. 2011). Overall, we estimate that the initial bottleneck is equivalent to that of a population of only 350 individuals, which is ∼24 times smaller than the initial number of French Canadian migrants to Quebec (Laberge et al. 2005). This initial bottleneck shared between core and front populations explains about 80% of the variance in allele frequencies at the expansion front, whereas only 20% of this variance can be attributed to the separate expansion of the ancestors of front individuals (**Fig. 5**). Note that this latter value should be considered as a lower bound for the total contribution of the expansion, because front and core samples have a shared history of being on the expansion front in the first few generations in Quebec, and this shared expansion is absorbed into the estimate of the bottleneck population size in our estimation procedure.

At first view, our estimations of selection coefficients (on the order of 10^−2^, **Fig. 5C**) for rare deleterious mutations are surprisingly higher than previous estimates (Eyre-Walker and Keightley 2007; Boyko et al. 2008; Racimo and Schraiber 2014; Henn et al. 2015b). A potential explanation for this apparent discrepancy is that our estimation is based on variants that were already rare (singletons) in Europe, and this set of variants should be enriched for more strongly deleterious variants than the set of all predicted deleterious mutations, which should include sites at high frequency (> 5%) that are presumably almost neutral (Boyko et al. 2008) despite being predicted as deleterious.

Overall, our results clearly suggest that due to the low effective size prevailing on the wave front of the colonization making selection less efficient than in the core, a small but significant mutation load has been generated in Quebec over a very short time (nine generations or less, see **Fig. 1** and Supplementary **Fig. S21**) by an increase in frequency of rare deleterious variants in front individuals by genetic drift. This excess of deleterious mutations on the front has probably only a minor effect on the total mutation load and on the fitness of most individuals, because these mutations are still at very low frequencies. Nevertheless, this wave front effect might be medically relevant as rare deleterious variants have a higher probability of being homozygous on the front than in the core, suggesting that rare recessive diseases should be more common in individuals whose ancestors lived on the front. In agreement with this prediction, we find that front individuals are indeed more likely to be derived homozygous for known pathogenic variants. Importantly, this effect is noticeably stronger than the relative risk to develop a rare disease because of inbreeding. In addition, the evidence of a relaxed selection on recent wave fronts suggests that prolonged periods of range expansions over hundreds of generations should have promoted the spread of deleterious mutations in newly settled territories, and have contributed significantly to global variation in mutation load and the burden of genetic diseases in modern humans.

## Methods

### Selection of individuals to sequence

We have selected individual to be sequenced by screening the genealogy of 3916 individuals of the CARTAGENE biobank (Awadalla et al. 2013), who could be connected to the BALSAC genealogical database (http://balsac.uqac.ca) thanks to the information they provided on their parents and grandparents. The BALSAC database includes records from all catholic marriages in Quebec from 1621 to 1965, totaling more than 3 million records (5 million individuals). The ascending genealogies of the 3916 CARTAGENE individuals were assessed for their maximum generation depth, their completeness defined as the fraction of ancestors that are traced back in an individual’s genealogy at generation *g* relative to the maximum number of ancestors (2^*g*^) at that generation (Jetté 1991), as well as our ability to assess the front or core status of the ancestors. We thus first eliminated 420 genealogies which spanned over less than 12 generations (maximum generation depth < 12 gen). We also eliminated 537 genealogies which had a mean depth smaller than 8 generations, 578 genealogies whose completeness (Jetté 1991) computed over the last 6 generations was less than 95%, and 97 additional genealogies whose completeness computed over the 12 generations was less than 30%. Genealogies were also filtered based on the quantity of information available for the computation of a cumulative Wave front Index (*cWFI*), defined as *CWFI* = Σ_*i*_ *GC_i_ × WFI_i_*, where the summation is over all ancestors in the genealogy, *GC_i_* is the genetic contribution of the *i*-th ancestor, *WFI_i_* is the wave front index of the *i*-th ancestor, defined as *WFI* = 1/(1 + *g*) (2011), and *g* is the number of generations elapsed since the foundation of the location where the ancestor reproduced (see ref. (Moreau et al. 2011) for more details). A *cWFI* value of 1 would imply that all the ancestors of the focal individual reproduced on the wave front. To ensure that differences in *cWFI* between individuals are not due to a lack of information on the core-front status of individuals in the genealogy, we eliminated 717 genealogies for which a single *WFI_i_* was missing for any individual of the 6 most recent generations (*WFI_i_* completeness <1 for the 6 most recent generations) and 15 additional genealogies for which the *WFI_i_* completeness until generation 12 was less than 0.5. We also excluded from the analysis genealogies for which the total number of individuals with computable *WFI* until generation 12 was either too small or too large, so that the *cWFI* was computed on genealogies of comparable total sizes. The 10% smallest and the 15% largest genealogies were thus eliminated (389 genealogies) from further analyses. The 1163 remaining individuals were ranked according to their *cWFI*, and we then selected individuals with the 10% smallest and 10% highest *cWFI*. We also eliminated from these two groups those individuals that were too closely related. The kinship coefficient *ϕ* (Wright 1922) was thus computed between all members of these groups to determine their relatedness. For 41 pairs of individuals more related than second cousins (*ϕ* > 1/64), one of the two individuals was removed at random. Finally, the 60 individuals with the lowest *cWFI* and the 60 individuals with the largest *cWFI* were selected for further DNA analyses. Among these, 51 individuals of each category for which peripheral blood samples were available in the CARTaGENE biobank were further considered for DNA extraction and sequencing. The geographic location of the marriage place of 102 individuals’ parents is reported in **Figure 1** and examples of the location of the ancestors of front and core individuals at various periods are shown in **Supplementary Animations S1** and **S2**.

### DNA extraction, library preparation and sequencing

Peripheral blood samples preserved in EDTA tubes from 102 selected individuals from the CARTaGENE cohort were processed for DNA extraction using the FlexiGene DNA kit as recommended by the supplier (Qiagen). Total DNA was quantified by measurements with the NanoDrop 8000 spectrophotometer (Themo Scientific) followed by dsDNA quantitation with the QUBIT 2.0 fluorometer (Life Technologies). DNA libraries were prepared for each sample following the standard protocol of KAPA Library Preparation Kit for Illumina sequencing platforms. A Covaris S2 fragmentation (Duty cycle - 10%, Intensity - 5.0, Cycle per burst - 200, Duration - 120 seconds, Mode Frequency - Sweeping, Displayed Power Covaris S2 – 23W) was performed on 1µg dsDNA input (50µl total volume) for each sample to generate 180 – 200 bp average size fragments. The resulting 3′ and 5′ overhangs were end repaired, 3’-adenylated and ligated to specific indexed adaptors. After a dual SPRI size selection of 250 – 450 bp adapter-ligated fragments, final pre-capture library enrichment was performed by LM-PCR followed by a library amplification cleanup with magnetic beads (AMpure XP, Agencourt). Following the protocol for whole exome capture with the Roche NimbleGen SeqCap EZ Exome + UTR Library kit (User’s Guide v4.2, http://www.nimblegen.com/products/seqcap/ez/exome-utr/index.html), the enriched fragments size distribution was then checked using a DNA 1000 chip on an Agilent 2100 Bioanalyzer for whole exome capture validation. The 102 uniquely indexed amplified DNA samples were mixed into 34 pool libraries of 3 different indexed DNA each, and were then hybridized to specific SeqCap EZ Hybridization Enhancing oligos at +47°C for 72 hours. After a washing step followed by a SeqCap EZ Pure Capture Beads recovery of the targeted sequences (here whole exome + UTRs), the multiplex DNA samples were amplified by a post-capture LM-PCR, cleaned with AMpure XP magnetic beads and bioanalyzed with a DNA 1000 chip to quantify and qualify the amplified captured multiplexed DNA samples. Prior to sequencing step, a final validation by qPCR assays was carried on the DNA samples to assess the relative fold enrichment in pre-captured sequences versus post-captured ones. Finally, these 34 DNA pools (one pool per lane) were paired-end (2x100bp) sequenced on an Illumina HiSeq 2500 System.

### Alignment and Variant Calling

Before mapping reads, a quality control was done using FASTQC, and trimming of the adapters and of poor quality read ends was done using Trim Galore (>=Q20). The reads were then mapped to the hg19 reference genome using BWA v 0.5.9r16 using the default parameters. PCR duplicates were removed using Picard-tools v1.56 (http://broadinstitute.github.io/picard/). We kept properly paired and uniquely mapped reads using Samtools v0.1.19-44428cd.

After these steps, we estimated the mean sequence coverage per individual, across the targeted exomic and UTR regions of cumulative length ∼106.5 Mb, to be between 67X-128X (**Supplementary Fig. S22**)

Realignment around indels and variants recalibration were performed with GATK v3.2-2. GATK v3.2-2 was also used to call variants using the workflow recommended by the Broad Institute (https://www.broadinstitute.org/gatk/guide/best-practices?bpm=DNAseq). We performed a first step using HaplotypeCaller, reporting the calls in GVCF mode. Then the joint genotyping calls were performed using the GenotypeGVCFs subprogram of GATK, to get the raw SNP and INDEL calls. The last step consisting in recalibrating and filtering the genotype calls was done with VQSR, using the recommended options separately on the SNP and INDEL calls.

### Sequence analysis

We removed all variants associated with a quality score below 30. We kept 426,301 SNPs and 43,081 indels and used ANNOVAR to functionally characterize these variants. **Supplementary Table S2** gives the number of variants in each ANNOVAR functional class.

Individual genotypes associated to low read depth (DP < 10) and low genotype quality (GQ < 20) were marked as missing genotypes.

We also collected polymorphism data for 305 individuals from 3 European populations (British from England and Scotland (GBR), Spanish from Spain (IBS) and Italians from Tuscany, Italy (TSI), **Supplementary Table S3**) from the 1000 Genomes phase 3 panel (The Genomes Project 2015). Note that the 1000 Genomes phase 3 panel set of variants consists of polymorphisms called from a combination of both low and high coverage data (between 8X - 30X). Our comparison of French Canadians and individuals from populations of the 1000 Genomes phase 3 panel was restricted to the genomic regions that were found in intersection between the targeted regions sequenced in the present study and the high coverage target of the 1000 Genomes phase 3 panel, which amount ∼46.4 Mb.

We defined shared SNPs between French Canadians and individuals from the 1000 Genomes phase 3 panel as SNPs found in both datasets.

Differences in number of various types of sites were obtained by a permutation test consisting in randomly permuting individuals between front and core, reestimating the desired statistics on the permuted samples and estimating the p-value of the observed statistics in the generated empirical null distribution.

### Assessment of mutation effects

The ancestral state of all mutations was characterized, following the 1000 Genomes project (The Genomes Project 2015), using the human ancestor genome inferred from the alignment of 6 primates (*Homo sapiens*, *Pan troglodytes*, *Gorilla gorilla*, *Pongo abelii*, *Macaca mulatta*, *Callithrix jacchus*) genomes (http://ftp.1000genomes.ebi.ac.uk/vol1/ftp/phase1/analysis_results/supporting/ancestral_alignments/) The biological impact of SNPs was assessed via GERP Rejected Substitution (GERP-RS) scores (Cooper et al. 2005; Davydov et al. 2010), which measure, at a given genomic location, the difference between the expected and the observed number of mutations occurring along a phylogeny of 35 mammals. GERP-RS scores were obtained from the UCSC genome browser (http://hgdownload.cse.ucsc.edu/gbdb/hg19/bbi/All_hg19_RS.bw). Note that the human sequence was not included in the calculation of GERP-RS scores. The human reference sequence was indeed excluded from the alignment for the calculation of both the neutral rate and site specific ‘observed’ rate for the RS score to prevent any bias in the estimates. Mutations were classified as being “neutral”, “moderate”, “large” or “extreme” for GERP-RS scores with ranges [-2,2[, [2,4[, [4,6[ and [6,∞[, respectively. GERP-RS scores of 0 indicates that the alignment of mammalian sequences was too shallow at that position to get a meaningful estimate of constraint (Goode et al. 2010) and sites with such scores were removed from all analyses involving GERP-RS scores.

We also used the CADD method (Kircher et al. 2014) to assess the functional effect of SNPs and to characterize short indels. CADD integrates many diverse annotations including conservation metrics, regulatory information, transcript information and protein-level scores into a single measure (C score) for each variant (Kircher et al. 2014). CADD has been implemented as a support vector machine and trained to differentiate human-derived alleles from simulated variants. The rationale for this choice is that deleterious variants are depleted by natural selection in existing but not simulated variation. We used scaled C-scores, phred-like scores ranging from 0.001 to 99, in our analyses, as these scores are easily interpretable. A scaled C-score larger than 10 indicates that the corresponding variant is predicted to be in the 10% most deleterious classes of variants. A scaled C-score larger than 20 indicates that the corresponding variant is predicted to be in the 1% most deleterious classes of variants. Mutations were classified as being “neutral”, “moderate”, “large” or “extreme” for CADD scores with ranges [0,10[, [10,20[, [20,30[ and [30,∞ [, respectively.

Most of our analysis in the main text relied on on SNPs and GERP-RS scores to assess their deleteriousness. We overall find very similar evidence of reduced selection in front populations using CADD scores for SNPs (**Supplementary Figs. S5,S7, S10 - S13**) or indels (**Supplementary Figs. S8 – S9**), suggesting that our results are robust to alternative deleteriousness scoring systems and to the choice of variants.

### Assessment of mutation load

Assess mutation load from genomic data is an inherently difficult problem (see e.g., (Lohmueller 2014) for a discussion of this problem). Instead, we use GERP-RS scores as a proxy for selection intensity and calculate, for each individual, the average GERP-RS score across all sites at which the focal individual carries a derived allele. We focus here on the average RS score per site. The average GERP-RS score per site is simply the average of GERP-RS scores calculated over all sites at which an individual carries at least one copy of a derived mutation: 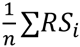, where n is the number of segregating sites per individual, and *RS_i_* is the GERP-RS score of site *i*. Note that this measure does not distinguish between heterozygous sites and derived homozygous sites. To account for the frequency of derived alleles we also calculated the average GERP-RS score across sites that have a given derived allele frequency.

### Detection of outlier SNPs and Gene Ontology analysis

To detect potential outlier SNPs based on levels of genetic differentiation, we used the outlier *F_ST_* method proposed by Beaumont and Nichols (Beaumont and Nichols 1996) and implemented in the Arlequin software (Excoffier and Lischer 2010). In brief, this test uses coalescent simulations to generate the joint distribution of *F_ST_* and heterozygosity between populations expected under a finite-island model, having the same average *F_ST_* value as that observed. This null distribution is then used to compute the *p*-value of each SNP based on its observed F_ST_ and heterozygosity levels. SNPs with *F_ST_* values outside the 99% quantile based on the simulations were considered as outliers. These SNPs were then annotated to Ensembl gene IDs with the R package BiomaRt (Durinck et al. 2009). SNPs were mapped to a gene if they were located in the gene transcript or within 10 kb to it. If a SNP was allocated to more than one gene with this method, we uniquely allocated to the gene to which it is closest. If more than one SNP was assigned to a given gene, we only kept the SNP with the highest *F_ST_* value.

We conducted a Gene Ontology (GO) enrichment analysis on the list of significant using the topGO R package (Alexa et al. 2006). We applied the default algorithm using a Kolmogorov-Smirnov (KS) test to detect highly differentiated biological processes and obtain their p-values. This approach integrates information about relationships between the GO terms and the different scores of the genes (here, the p-values) into the calculation of the statistical significance. We kept in this analysis only GO terms which included more than 10 genes.

### Maximum likelihood estimation of past demography and selection coefficients

We considered sites that are found as private singletons in the European 1000G populations and that are found polymorphic in Quebec. We used the current frequency of these variants in Europe as a proxy for their frequency during the foundation of Quebec. This allows us to directly estimate front and core effective population sizes without having to estimate additional parameters for the European population.

We modeled the evolution of allele frequencies at independent sites under random genetic drift and natural selection in two panmictic populations, denoted the core and the front. Variables describing properties of the front and core are denoted with sub- or super-scripts *f* and *c*, respectively. For simplicity, we only present calculations for the front. The core can be treated analogously. Then 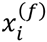 denotes the number of sites with a derived allele frequency of *i*. Let 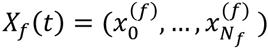, denote the SFS on the front where *N_f_* is the effective population size at the front and *t* denotes the time (in generations) since the founding of Quebec. Assuming a Wright-Fisher model of drift and genic selection (that is, no dominance or epistasis), the SFS then evolve according to

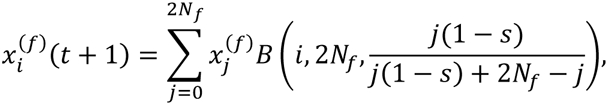

where *B*(*k, n, p*) denotes the binomial distribution and *s* is the strength of selection against the derived allele. We calculate the current allele frequency distribution (16 generations after the onset of the settlement) *X_f_* (16) with the initial condition 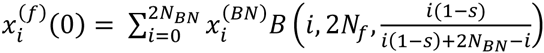, where 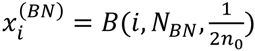 is the expected allele frequency distribution in the bottlenecked population and *n*_0_ is the sample size in Europe. We then obtain the expected allele frequency distribution for a sample of *n_f_* = 51 individuals by

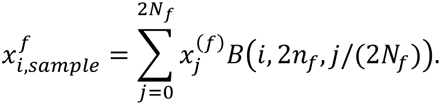

Let 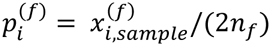 be the relative frequency of sites with a derived allele frequency of *i*. To account for the fact that we only consider sites shared between Europe and Quebec, we correct the allele frequency distribution by multiplying the proportion of sites that are not found polymorphic at the front, 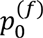, by 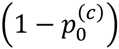, i.e., we count only the proportion of sites where the derived allele is lost in the front but that are polymorphic in the core, and then renormalize such that 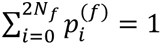. We can then calculate the likelihood from our data as

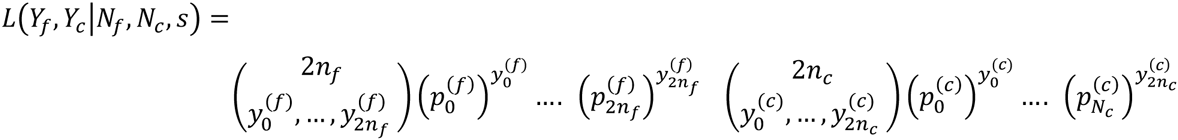

where 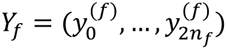 and 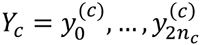 denote the observed derived allele frequencies in front and core respectively. The likelihood was then maximized numerically via a grid search in the parameter space.

### Individual Based Simulations

We performed individual based simulations of a range expansion in a 2D habitat consisting in a lattice of 11x11 discrete demes (stepping stone model). Generations are discrete and non-overlapping, and mating within each deme is random. Migration is homogeneous and isotropic, except that the boundaries of the habitat are reflecting, i.e., individuals cannot migrate out of the habitat. Population size grows logistically within demes. Our simulations start from a single panmictic ancestral population, representing France. After a burn-in phase that ensures that the ancestral population are at mutation-selection-drift balance, a propagule of founders is placed on the deme with coordinates (3,6) on the 11x11 grid representing French Canada (see **Supplementary Fig. S16)**. During the next 6 generations, the population expands along a 1 deme wide corridor in the middle of the habitat (representing the St-Laurent river corridor). During these 6 generations, all colonized demes in French Canada receive migrants from the ancestral populations in equal proportions. The number of migrants were chosen to roughly match historical records (Haines and Steckel 2000). In particular, we chose 1000, 2000, 1000, 1000, 1000, and 2000 pioneer immigrants from the ancestral population for the first 6 generations, respectively. After that, the expansion continues into the remaining habitat for 11 generations. See **Supplementary Fig. S16** for a sketch of the model.

We chose a carrying capacity of *K* = 1,000 diploid individuals and the size of the ancestral population was 10,000. Migration rate was set to *m* = 0.2 and the within deme growth rate was *R* = 2 (that is, at low densities the population doubles within one generation, reflecting the average absolute fitness of approximately 4 – 5 surviving children getting married per women (Moreau et al. 2011)). We simulated a set of 10,000 independent biallelic loci per individual. The genome-wide mutation rate was set to *u* = 0.1. Mutations occur only in one direction and back mutations are ignored. We performed two types of simulations: (i) evolution of neutral mutations, and (ii) evolution of sites under purifying selection. In the latter case, we assumed that all sites had the same selection coefficient *s*. Mutations interact multiplicatively across and within loci, that is, there is no dominance or epistasis. We also simulated and recorded the cumulative wave front index (cWFI) of each individual. The simulation code can be downloaded from: https://github.com/CMPG/ADMRE.

## Data Access

Requests for data published here should be submitted to the corresponding authors, citing this study.

## Acknowledgements

We are grateful to Claude Bhérer for her detailed comments on the manuscript. We would like to thank the CARTaGENE participants and team for data collection and assistance, Marc Tremblay for his help in connecting CARTaGENE individuals to the Balsac genealogical data base, Remy Brugmann for Bioinformatic analyses, and the Ubelix High Performance Computing cluster of the University of Bern. We confirm that informed consent was obtained from all subjects. This work has been made possible by a Swiss NSF grant No. 31003A-143393 to LE. AH is an FRSQ Research Fellow. AH currently holds a Career Development Fellowship as part of the eMedLab Medical bioinformatics partnership funded by the Medical Research Council, UK. PA is supported by the Ministry of Research of Ontario.

